# Molecular characterization of a DyP-type peroxidase from the human parasitic cestode *Echinococcus multilocularis*

**DOI:** 10.1101/2022.11.07.515413

**Authors:** Johannes Ulrich, Klaus Brehm

## Abstract

The lethal zoonosis alveolar echinococcosis is caused by the metacestode larval stage of the tapeworm *Echinococcus multilocularis*. During the chronic phase of the disease, metacestode tissue is growing infiltratively into liver tissue and provokes an immunes response of the host. Mechanisms of parasite defence against reactive oxygen species (ROS), which are produced during parasite growth and host immune responses, are incompletely understood so far. We herein describe the characterization of an *Echinococcus* Dyp (dye decolorizing) – type peroxidase, EmDyp, family members of which are typically expressed by bacteria and fungi. EmDyp showed significant homologies to bacterial and fungal Dyp peroxidases and recombinantly expressed EmDyp displayed profound enzymatic activity towards different substrates such as 3,3’-diaminobenzidine or luminol. Furthermore, although structurally not being related to classical catalases, EmDyp showed catalase activity in respective activity gels. *In situ* hybridization experiments showed expression of the EmDyp expressing gene, *emdyp*, in the germinal layer of the metacestode as well as in the posterior region of the protoscolex, both in differentiated and in germinative (stem) cells of the parasite. Interestingly, RT-qPCR experiments demonstrated that *emdyp* expression is induced in the metacestode upon growth under aerobic conditions. Particularly high expression of *emdyp* was observed under *in vivo* growth conditions in jirds within the liver. These data indicate a role of EmDyp in the defence of the metacestode against host- and/or parasite-derived ROS during chronic alveolar echinococcosis. Since Dyp-type peroxidases are not encoded on the genomes of mammalian hosts for *E. multilocularis*, EmDyp might be used as a target molecule for developing novel therapeutics against the parasite.

## Introduction

The metacestode larval stage of the fox-tapeworm *Echinococcus multilocularis* is the causative agent of alveolar echinococcosis (AE), a lethal zoonosis prevalent in the Northern Hemisphere (Romig et al., 2017). After initial infection of the intermediate host (rodents, humans) by the oncosphere, the parasite undergoes a metamorphosis-like transition towards the cystic metacestode stage within the host liver (Brehm and Koziol, 2017). The metacestode subsequently growth infiltratively, like a malignant tumour, within the host organs eventually leading to malfunctions and organ failure (Brunetti et al., 2010). Metacestode growth is exclusively driven by a population of pluripotent stem cells, the so-called germinative cells, which account for 25% of all cells of the metacestode germinative layer (Koziol et al., 2014). At the end of an infection of natural intermediate hosts (rodents), numerous brood capsules are formed by the germinative cells of the metacestode, giving rise to protoscoleces, which are the infective form for the definitive host (foxes, dogs) (Brehm and Koziol, 2017).

Previous immunological studies indicated an important role of macrophages and neutrophils in host defence against *Echinococcus*, particularly during the early phase of parasite establishment within the intermediate host (Alkarmi and Behbehani, 1989; Zhang et al., 2003; Wang et al., 2020). Due to the decisive role of reactive oxygen species (ROS) in effector mechanisms of macrophages and neutrophils (Silva, 2010), it is thus reasonable to assume that protection against ROS is important in *Echinococcus* establishment and intrahepatic growth. Although the precise energy metabolism pathways used by *E. multilocularis* during *in vivo* growth within the liver of the intermediate host have not yet been fully elucidated, the relative oxygen tension in this compartment (~ 5%; Guo et al., 2017; Chaudhry et al., 2022, Martinez-Gonzalez et al., 2022) indicates that the parasite at least to a certain extent uses classical, oxygen-consuming mechanisms which necessarily result in the production of ROS such as superoxide anion and hydrogen peroxide. Hence, for both protection against the host immune response and DNA protection during aerobic growth, *E. multilocularis* must be equipped with appropriate detoxification systems to counteract the activities of ROS.

Within the genus *Echinococcus*, ROS detoxification systems have so far mostly been studied in the closely related species *E. granulosus*, who causes infections in domestic animals. Very recently, Cancela et al. (2019) identified several proteins that are upregulated in *E. granulosus* protoscoleces upon incubation with exogenous hydrogen peroxide, and many of those have presumed functions in ROS detoxification. One of the best studied systems in this regard is a thioredoxin-glutathione system that is expressed in both the cytosolic and the mitochondrial department of *E. granulosus* cells (Salinas et al., 2004; Bonilla et al., 2008). Furthermore, members of the glutathione-S-transferase family have been reported in *E. granulosus* (Fernandez et al., 2000; Iriarte et al., 2012) and at least within the genome of both *E. multilocularis* and *E. granulosus*, superoxide dismutases are encoded (Tsai et al., 2013). Genes encoding classical catalases are, however, absent in cestode genomes (Tsai et al., 2013).

Dye-decolorizing peroxidases (DyP) have originally been described in bacteria and fungi and constitute a class of heme peroxidases which structurally differ from classical peroxidases (Sugano, 2009; Zamocky et al., 2015; Sugano and Yoshida, 2021). As other heme peroxidases, Dyp-type peroxidases reduce hydrogen peroxide and then use the resulting electrons for oxidizing a variety of different substrates, amongst which are also numerous synthetic dyes (Colpa et al., 2013). Structurally, Dyp-type peroxidases differ from other peroxide classes in that they possess a highly conserved GXXDG motif as part of the heme binding site and that they form two antiparallel ß-sheets of which one participates in heme binding (Sugano, 2009; Zamocky et al., 2015; Sugano and Yoshida, 2021). In both bacteria and fungi, secreted and intracytoplasmic forms of Dy-type peroxidases exist, and it is assumed that they participate in protecting cells from endogenous oxidative stress, produced during aerobic metabolism, as well as from exogenous ROS (Sugano and Yoshida, 2021). Furthermore, Dyp-type peroxidases also exhibit a relatively broad pH spectrum and are also active at low pH (Zamocky et al., 2015; Sugano and Yoshida, 2021). Due to their dye-decolorizing activities, Dyp-type peroxidases have so far mostly been studied in an industrial context as biodegradative enzymes of chemical waste (Colpa et al., 2013; Sugano and Yoshida, 2021), but in the case of mycobacterial isoforms, a role in host defence against ROS produced during the immune response has been suggested (Kirykowicz and Woodward, 2020). Interestingly, Dyp-type peroxidases are not confined to bacteria and fungi but have recently also been found to be encoded by the genomes of invertebrates and it has been hypothesized that they were acquired by horizontal gene transfer (Zamocky et al., 2015). As yet, no bilaterian member of the Dyp-type peroxidase family has been biochemically characterized, nor have functional investigations been carried out towards the role of these enzymes in parasite biology. In the present work we show that *E. multilocularis* expresses a functionally active Dyp-type peroxidase and that the respective gene is highly induced in parasite tissue upon growth within host tissue. A role of the enzyme in protecting parasite cells from ROS that are produced during aerobic growth and during host responses is discussed.

## Materials and Methods

### Ethics statement

*In vivo* passage of parasite tissue was performed in mongolian jirds (*Meriones unguiculatus*), which were raised and housed at the animal facility of the Institute of Hygiene and Microbiology, University of Würzburg. This study was performed in strict accordance with German (*Deutsches Tierschutzgesetz,TierSchG*, version from Dec-9-2010) and European (European directive 2010/63/EU) regulations on the protection of animals. The protocol was approved by the Ethics Committee of the Government of Lower Franconia (Regierung von Unterfranken) under permit number 55.2.2-25322-1479-8.

### Organisms and culture methods

Experiments involving metacestode vesicles were performed with the *E. multilocularis* isolate H95 which derived from a naturally infected fox of the region of the Swabian Mountains, Germany (Jura et al., 1996). Isolate H95 has lost the capacity to produce brood capsules and protoscoleces and exclusively grows as metacestode tissue both *in vitro* and *in vivo*. Experiments involving protoscoleces were performed using isolates SUMA16 and RD15 which derive from Old World Monkey species (*Macaca fascicularis*) that had been naturally infected in a breeding enclosure in the region of Göttingen, Germany (Tappe et al., 2007). SUMA16 and RD15 are recent isolates that are fully capable of protoscolex production *in vitro* and *in vivo*. Metacestode vesicles were either cultivated *in vitro* in the presence of host feeder cells (rat Rheuber hepatoma) or under axenic conditions (nitrogen atmosphere, reducing agents) as previously described (Spiliotis and Brehm, 2009). *Echinococus* primary cell cultures were established and maintained in culture as described by Spiliotis et al. (2008). Culture media were changed every three to four days.

### PCR amplification and cloning procedures

Single-stranded cDNA was produced from mRNA isolated from in vitro cultivated metacestode vesicles essentially as previously described (Förster et al., 2019) and was used as a template in all PCR and cloning procedures. For the amplification of full-length *emdyp* transcripts, the primers ipoxGfw2 (5’-GAT GAG TTA CTC GAG TCG C-3’) and ipoxR2G (5’-GTT ACA GCA GGC TAA AAC G-3’) were used (amplifying the shorter reading frame) as well as ipox5PR (5’-GTT CGA TGA GAG GTT TTG ATT TGC-3’) and ipoxR2G (amplifying the longer transcript). PCR products were then blunt end cloned into vector pJET2.1 (Thermo Fisher Scientific) and sequenced from both sides. For heterologous expression in *E. coli*, the *emdyp* reading frame was cloned into expression plasmid Pet151 D (Thermo Fisher) using the AQUA cloning procedure as described by Beyer et al. (2015). To this end, primers xipoxf (5’-GAA AAC CTG TAT TTT CAG GGT AGT TAC TCG AGT CGC CTG ACA GA-3’) and xipoxr (5’-GAA ATC AGT TTC TGT TCT AGT TAC AGC AGG CTA AAA CGC TTT CCA-3’) were used for PCR amplification of the *emdyp* open reading frame and cloned into Eco RI digested Pet151 D, thus leading to a fusion of the *emdyp* open reading frame with the anti-V5 epitope tag (Invitrogen) and a hexahistidine tag. The resulting plasmid was named Pet151-DyP-V5.

### Induction and purification of recombinant EmDyp

*E. coli* strain BL21 (DE3) (New England Biolabs) was transformed with Pet151-Dyp-V5 and incubated over night at 37°C. Induction of recombinant protein expression was subsequently carried out using 0,12 mM isopropyl-ß-D-thiogalactopyranosid (IPTG) essentially as previously described (Liu et al., 2011). Purification of recombinant EmDyp was subsequently carried out according to Liu et al. (2011) with several slight modifications. Briefly, thawed bacterial pellets were taken up in lysis buffer (25 mM Tris-HCl, pH 8; 200 mM NaCl; 1 mM PMSF (solved in DMSO)) and lysis was carried out for 60 sec using ultrasonication. 10 ml lysis suspension were then incubated for 30 min at RT with 20 μl heme solution (100 mg heme (Sigma Aldrich) dissolved in 6 ml NaOH), resulting in a colour change towards red. The resulting suspension was centrifuged for 20 min at 28.000 g and the supernatant was incubated over night at 4°C with ProBond resin (Invitrogen). Probes were then centrifuged for 5 min at 500 g, the pellet was solved in lysis buffer and transferred onto a PD-10 desalting column (GE Healthcare). Purification was then carried out essentially according to the manufacturer’s protocol using elution buffer (lysis buffer with 250 mM imidazole) for final elusion.

### Acrylamide gel electrophoresis, Western blot detection, and catalase activity gels

12% SDS-PAGE and Western blot detection was carried out essentially as previously described (Spiliotis et al., 2006) using an alkaline phosphatase-coupled, monoclonal murine antibody directed against the V5 epitope (anti-V5-AP; Invitrogen) of recombinant EmDyp. *In situ* catalase activity in native polyacrylamide gels was measured essentially as previously described (Haas et al., 1999), omitting staining for peroxidase activity and using exclusively hydrogen peroxide for detecting enzymatic activities.

### Peroxidase activity assays

For measuring peroxidase activity towards 3,3’-diaminobenzidine (DAB) as a substrate, previously described methodology (Herzog and Fahimi, 1973) using 0,1 % gelatine solution was employed. Briefly, 30 μl of substrate buffer (0,1% gelatin; 0.13 mg/ml DAB tetrahydrochloride; 25 mM Tris-HCl, pH 8; 200 mM NaCl) were mixed with different amounts of recombinant EmDyp (up to 30 μg) and 20 μl hydrogen peroxide solution (30%). Absorption at 465 nm was then measured spectrophotometrically for up to 5 min. For measuring EmDyp activities towards luminol, the Pierce ECL Western blot substrate kit (Thermo Fisher) was used essentially according to the manufacturer’s instructions. All experiments were carried out in triplicates.

### Whole mount in situ hybridization (WISH) and EdU labelling

WISH was carried out on *in vitro* cultivated metacestode vesicles and on activated protoscoleces essentially as previously described (Koziol et al., 2014; Koziol et al., 2016) after a 5 h pulse of incorporation of 5-ethynyl-2’-deoxyuridine (EdU) for staining proliferative cells (Koziol et al., 2014). *emdyp* WISH probe preparation was carried out essentially as described (Koziol et al., 2014) using primers ipoxwish (5’-GAT GAA GAC AAC GAG TTG ATA G −3’) and ipoxR2G (see above) for amplification from *E. multilocularis* metacestode cDNA. *In vitro* cultivation of metacestode vesicles in the presence of host feeder cells and under axenic conditions were performed as previously described (Spiliotis and Brehm, 2009) and the activation of protoscoleces by low pH, pepsin treatment was performed as described in Gelmedin et al. (2010).

### Quantitative real-time RT-PCR (RT-qPCR)

For qPCR, cDNA was first synthesized from different samples (metacestode vesicles under aerobic and anaerobic growth conditions, activated and dormant protoscoleces, in vivo parasite material from peritoneum and liver infiltrate) as previously described (Koziol et al., 2014) using 100 ng of total RNA as input. qPCR was then carried out using a StepOne Plus Realtime PCR cycler (Applied Biosystems) and a PCR mix of 1 x HOT FIREPol EvaGreen qPCR Mix containing 300 nM of each primer and cDNA according to previously established protocols (Perez et al., 2019; Ancarola et al., 2020). emdyp specific PCR was performed using primers ipoxqpcr-f (5’-ATT CGT CGA TGG CCA GCA-3’) and ipoxqpcr-r (5’-GGA GGC AAT CTT CTT CAT GTC G −3’) and the constitutively expressed gene *elp* (EmuJ_000485800; Ref) was used as a control using previously established primers (Ancarola et al., 2020). Cycling conditions were 15 min at 95°C, followed by 40 cycles of 15 sec at 95°C, 20 sec of 58°C, and 20 sec of 72°C. Data collection was performed at 72°C. PCR efficiencies were using LinRegPCR (Untergasser et al., 2021), amplification product specificity was assessed by melting curve analysis and gel electrophoresis. Expression levels were calculated by the efficiency correction method using cycle threshold (Ct) values according to Ancarola et al. (2020).

### In silico analyses and statistics

*E. multilocularis* genome mining and BLASTP analyses have been carried out on WormBase ParaSite (Howe et al., 2016) using Annotation version 2015-12-WormBase for *E. multilocularis* and 2022-01-Worrmbase for *Schistosoma mansoni* (https://parasite.wormbase.org/index.html). Reciprocal BLASTP analyses against SWISSPROT database have been performed on GenomeNet (https://www.genome.jp/). Multiple sequence alignments were performed using Clustal Omega (Madeira et al., 2022) with default settings as offered on (https://www.ebi.ac.uk/Tools/msa/clustalo/). Protein domain analyses were performed using SMART (http://smart.embl-heidelberg.de/) (Letunic et al., 2021). Statistical analyses were performed using JASP (https://jasp-stats.org/) software, version 0.16.3.0, employing classical one-way ANOVA.

## Results and Discussion

### Characterization of the E. multilocularis Dyp-type peroxidase encoding gene

The *E. multilocularis* whole genome project revealed a gene model, EmuJ_001133900, the product of which had been annotated as iron-dependent peroxidase (Tsai et al., 2013). Since the *E. multilocularis* metacestode most likely encounters ROS during an infection of the intermediate host, and since no peroxidases had so far been described in this organism, we were interested in a closer characterization of EmuJ_001133900. By inspecting metacestode and protoscolex transcriptome data, which had been collected during the genome sequencing project (Tsai et al., 2013), on WormBase Parasite (https://parasite.wormbase.org/index.html), we obtained clear indications that EmuJ_001133900 is alternatively spliced at the 5’ end, leading to different transcripts that can be distinguished by RT-PCR (Fig. 1). Based on *E. multilocularis* genome information, we therefore designed different primers for the amplification of both transcript versions, henceforth designated as transcript A, using a splice donor site close to the transcript 5’ end (SD1), and transcript B, using a splice site 88 bp downstream of SD1 (SD2). Both versions were then PCR amplified from protoscolex cDNA and fully sequenced, thus verifying the predictions made during the genome sequencing project. As shown in Fig. 1, both splice sites employed canonical GT dinucleotides at the donor end and both used the same splice acceptor site (SA), 450 nt downstream of SD2.

**Fig. 1.:**
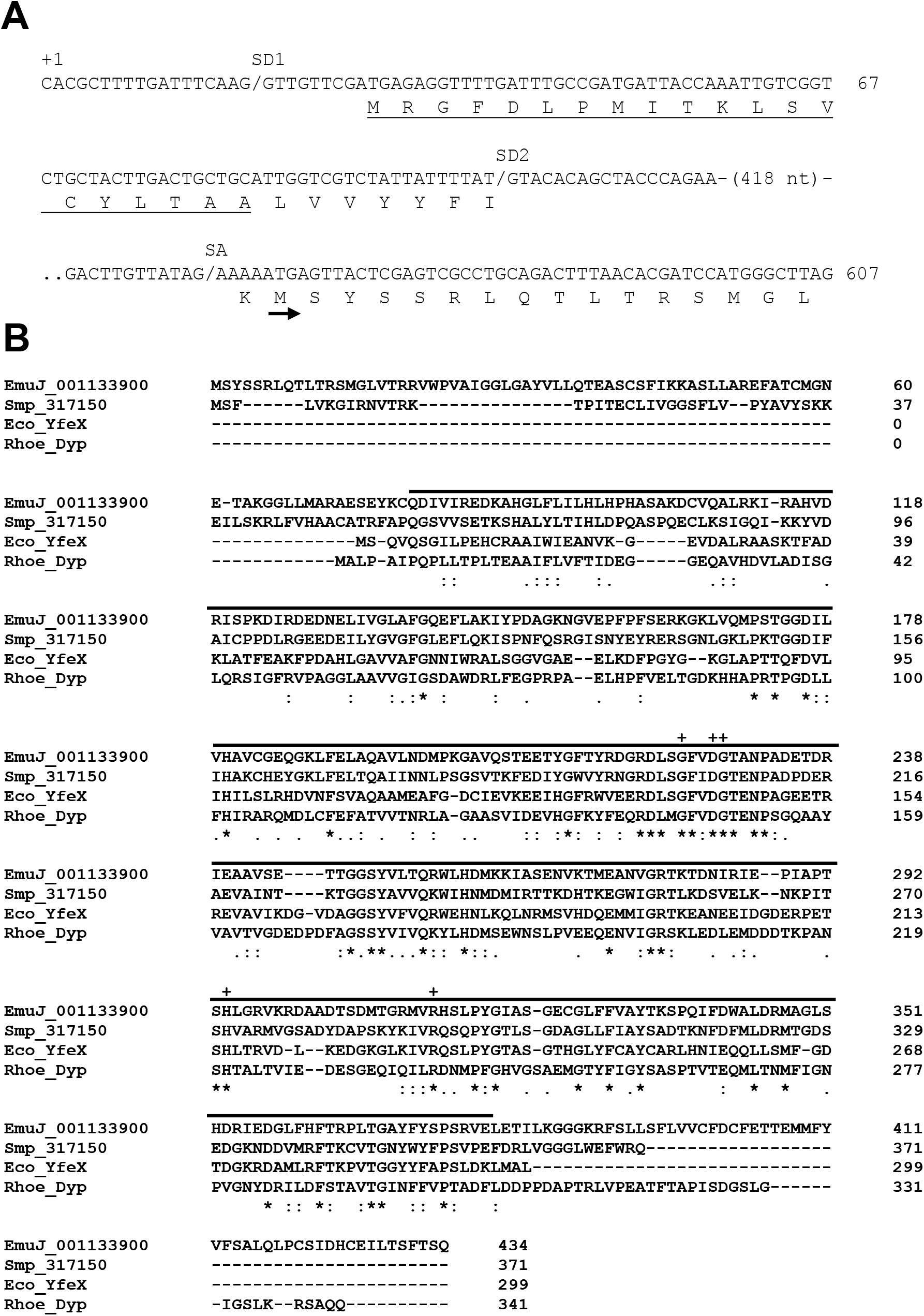
*emdyp* cloning and sequence features. (A) Alternative 5’ end splicing of *emdyp*. Depicted are the *emdyp* cDNA sequence at the transcript 5’ end. ‘+1’ indicates the transcription start site. The deduced amino acid sequence is shown below the nucleotide sequence. ‘SD1’ and ‘SD2’ indicate alternative splice donors, ‘SA’ the splice acceptor used by both donors. A small arrow indicates the start methionine used for translation of the cytoplasmic version of EmDyp. The putative signal peptide of the secreted version is underlined. Numbers to the right indicate the distance of the primary transcript to the transcription start site. (B) Sequence homologies of EmDyp. Shown is an alignment of EmDyp (EmuJ_001133900), a putative Dyp-type peroxidase homolog from *Schistosoma mansoni* (Smp_317150) and two bacterial Dyp-type peroxidases from *E. coli* (Eco_YfeX) and *Rhodococcus equi* (Rhoe_Dyp). The Dyp-type peroxidase domain as identified by SMART analysis is marked by a black line above the alignment. Sites of perfect alignment (*) as well as those of strong (:) or weak (.) similarity are marked below the alignment. Residues highly conserved in all Dy-type peroxidases are marked by ‘+’ above the alignment. UniProt accession codes for the aligned sequences are A0A068YJZ2 (EmDyp), A0A5K4F6L3 (Smp_317150), Q8XBI9 (Eco_YfeX), and C0ZVK5 (Rhoe_Dyp).

We then analysed the deduced amino acid sequences of transcript A (total length 434 amino acids) and B (461 amino acids) for domain structures and in both cases identified a predicted Dyp-type peroxidase domain of 300 amino acids (Fig. 1). Interestingly, for the longer protein version encoded by transcript B, a transmembrane helix was predicted close to the N-terminus (Fig. 1), which highly likely serves as an N-terminal signal peptide for protein export. The smaller EmDyp version encoded by transcript A, on the other hand, did not contain an N-terminal signal peptide. It thus appears that the two transcript versions of *emdyp* encode intracytoplasmic (transcript A) and secreted (transcript B) isoforms of the same enzyme.

When we performed BLASTP analyses on the SwissProt database, highest homologies were obtained for both versions of EmDyp with the well-characterized Dyp-type peroxidase YfeX of *Escherichia coli* (Dailey et al., 2011) as well as with several other bacterial Dyp-type peroxidases (Figure 1). Although overall sequence homologies were quite moderate (e.g. 36% identical and 51% similar residues to YfeX), EmDyp contained all amino acid residues important for heme binding and enzyme function at the corresponding sequence positions (Fig. 1). Hallmarks of plant peroxidases such as a conserved R-x-x-F/W-H motif (Sugano, 2009), for example, were missing in EmDyp. Furthermore, the parasite enzyme was missing clear homologies to classical catalases. We thus concluded that EmuJ_001133900 encodes a typical Dyp-type peroxidase and named the gene *emdyp*. Interestingly, when we performed BLASTP analyses against the genome of the related parasite *Schistosoma mansoni* in WormBaseParaSite, we identified a gene encoding a Dyp-type peroxidase (Smp_317150) which was highly homologous to the *emdyp*-encoded protein, EmDyp (Fig. 1). We thus concluded that Dyp-type peroxidases are also expressed by trematodes.

Among bacterial Dyp-type peroxidases, both intracellular and extracellular (within the periplasmic space) forms exist, and it is assumed that the intracellular peroxidases are mostly involved in the removal of ROS that are produced during aerobic growth (Sugano and Yoshida, 2021). One presumed role of the periplasmic peroxidases of bacteria and several secreted fungal Dyp-type peroxidases, on the other hand, is the protection of cells from oxidative stress exerted by host immune cells. Since *emdyp* obviously encodes intracellular and secreted forms of EmDyp, it is thus interesting to speculate that the parasite protein is also involved in both, protection from intracellular ROS produced during aerobic growth, and defence against immune effector cells. Most interestingly, a similar situation to that of *emdyp* has already been observed for the thioredoxin-encoding gene *egtgr* of the related cestode *E. granulosus*, by which cytosolic and mitochondrial enzyme variants are produced involving alternative splicing from an identical primary transcript (Chalar et al., 1999; Agorio et al., 2003). In the case of *egtgr*, alternative 5’ splice acceptor sites within the primary transcript are utilized, which acquire the canonical *Echinococcus* spliced leader exon during trans-splicing (Brehm et al., 2000), and which code for longer (mitochondrial) or shorter (cytosolic) polypeptides (Agorio et al., 2003). When we performed RT-PCR experiments on *emdyp* using primers against the *Echinococcus* spliced leader (Brehm et al., 2000), no indications were found that the peroxidase encoding transcripts are trans-spliced (data not shown). Nevertheless, the utilization of alternative splicing at primary transcript 5’ ends for producing protein variants of different cellular location seems to be more frequently used by *Echinococcus*.

### Enzymatic activities of EmDyp

We next investigated enzymatic activities of EmDyp. To this end, the EmDyp reading frame encoding the cytosolic version (i.e. excluding any signal peptide sequence) was cloned into an expression plasmid as a fusion protein to the V5 antibody epitope and was heterologously expressed in *E. coli*. The fusion protein was then purified involving incubation with hemin upon which colour changes clearly indicated that EmDyp is a heme binding enzyme. The purified protein was then used in assays for peroxidase activity using different substrates. As shown in Fig. 2A, when we used DAB, a common substrate for Dyp-type peroxidases (Sugano, 2009; Sugano and Yoshida, 2021), EmDyp displayed clear peroxidase activity depending on the amount of enzyme added. The assay was also carried out at different pH values and as shown in Fig. 2B, EmDyp displayed relatively high activities over a range from pH 4 to pH 8 as it is typical for Dyp-type peroxidases (Sugano, 2009; Sugano and Yoshida, 2021). In another enzymatic assay, we clearly detected oxidising activity of purified EmDyp towards luminol (Fig. 2C). On the other hand, when we used Remazol brilliant blue (RB19) as a substrate for EmDyp, no enzymatic activity was observed (data not shown).

**Fig. 2:**
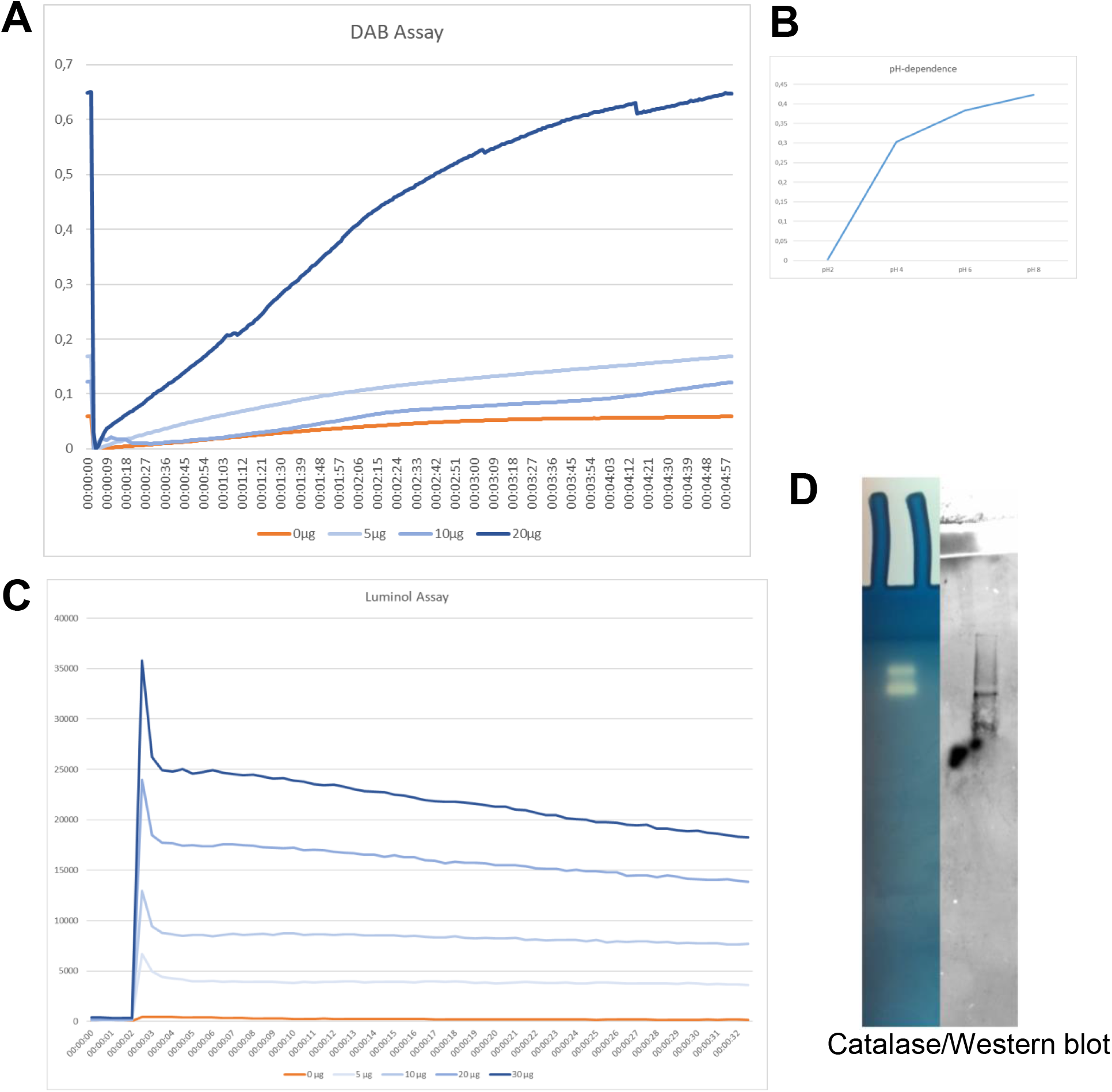
EmDyp catalytic activities. (A) Activity of purified recombinant EmDyp using DAB as a substrate. Curves indicate OD_465nm_ upon addition of different amounts of purified EmDyp over a reaction time of 5 min. DAB was used as a substrate. (B) pH dependence of EmDyp activity towards DAB. The assay was carried out as in (A), but at different pH values, as indicated. (C) Activity of purified recombinant EmDyp towards luminol. Curves indicate activity units upon addition of different amounts of purified EmDyp over a reaction time of 30 sec. (D) Catalase activity of EmDyp. 30 μg of recombinantly expressed and purified EmDyp were run on a native PAA gel and stained for catalase activity (left gel) according to (Haas et al., 1991). The proteins were also blotted onto a membrane and Western blot was carried out using an anti-V5 antibody (right-hand side).

During our assays using DAB as a substrate, we observed the formation of gas, indicating that EmDyp might also act as a catalase. To closer investigate this aspect, we performed native gel electrophoresis on the purified parasite enzyme followed by staining for catalase activity. As shown in Fig. 3, we could detect catalase activity at two different bands, which corresponded to EmDyp as also assessed by Western blotting directed against the V5-epoitope of the recombinant enzyme. Hence, unlike several other Dyp-type peroxidases (Chen et al., 2015), EmDyp appears to also possess catalase activity.

**Fig. 3:**
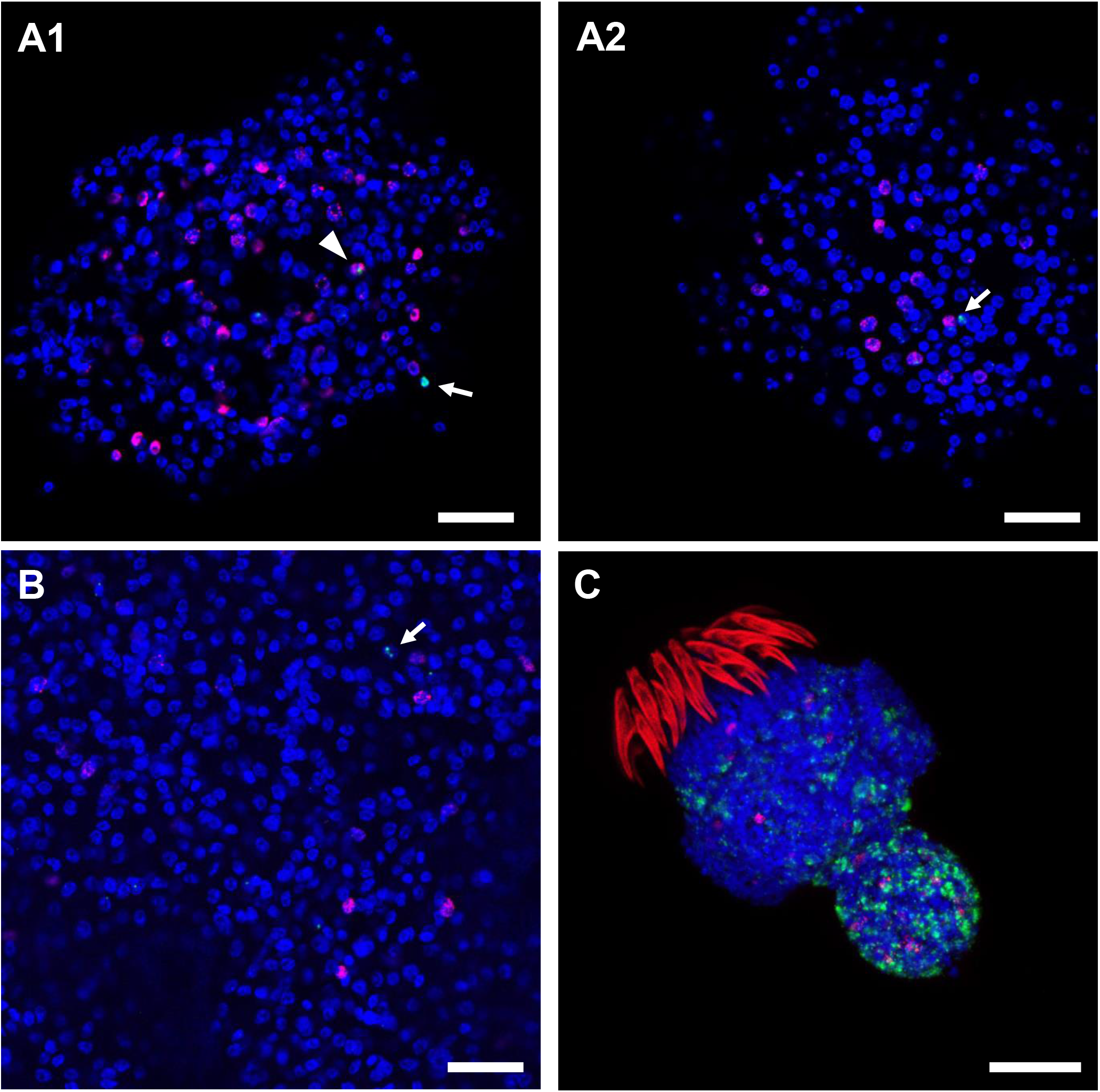
WISH for expression of *emdyp* in *Echinococcus* larval stages. Double staining for EdU-uptake (stem cells; red) and WISH using an anti-*emdyp* probe (green) has been carried out on *Echinococcus* primary cells (A1, A2), metacestode vesicles (B), and activated protoscoleces (C). DAPI staining (blue) has been performed to visualize nuclei. White arrows indicate cells positive for *emdyp* but negative for EdU staining. A white triangle indicates EdU+/*emdyp*+ cell. White bar represents 20 μm in A and B, 50 μm in C.

Taken together, our assays so far clearly indicated that EmDyp is an active enzyme and can act as peroxidase and catalase. As such, EmDyp could fulfil an important detoxifying role against ROS during *Echinococus* growth within the host.

### Expression of emdyp in Echinococcus larval stages

We next investigated the expression of *emdyp* in different parasite larval stages, employing a previously established protocol for WISH (Koziol et al., 2014). To assess whether *emdyp* is expressed in *Echinococcus* germinative (stem) cells, we combined WISH with EdU-staining, thus identifying stem cells that are in the S phase of the cell cycle (Koziol et al., 2014). First, we carried out combined EdU/WISH staining on *Echinococcus* primary cell cultures and identified only few cells positive for *emdyp* expression (Fig. 3A). Of these, the majority did not show co-staining with EdU, although in few instances EdU+/*emdyp*+ cells were identified (Fig. 3A1). We thus concluded that *emdyp* is expressed in both stem cells and postmitotic cells, albeit to a very low level in primary cell cultures. Second, we investigated the germinative layer of *in vitro* cultivated metacestode vesicles. As shown in Fig. 3B, very few cells expressed *emdyp*. Again, both EdU+ and EdU-cells were seen to be positive for *emdyp* (data not shown). Finally, we carried out WISH on activated protoscoleces and found intense *emdyp* signals, particularly for the posterior part of the scolex, although fewer signals were also observed in the sucker region (Fig. 3C). Again, the majority of *emdyp+* cells were EdU negative, but few were positive for both signals. Taken together, our WISH analyses indicated strong expression of *emdyp* in the protoscolex and only very few *emdyp*+ cells in primary cell cultures and the metacestode.

We next investigated *emdyp* expression in the metacestode and in the protoscolex by RT-qPCR in comparison to the constitutively expressed houskeeping gene *elp* (Brehm et al., 1999). To this end, we prepared cDNA from mRNA isolated from non-activated (dormant) protoscoleces, from protoscoleces activated by low pH/pepsin treatment, from axenically cultivated metacestode vesicles, kept in the absence of oxygen, and from metacestode vesicles cultivated in the presence of host feeder cells and in the presence of oxygen. Furthermore, we isolated mRNA from metacestode material, which did not contain brood capsules or protoscoleces, from the peritoneum and from the liver of infected jirds. As shown in Fig. 4A, protoscoleces displayed a markedly higher (> 5.000-fold) expression of *emdyp* than axenically cultivated metacestode vesicles. Activation of protoscoleces by low pH/pepsin, which mimics the passage through the digestive tract of the definitive host, led to a slight increase of *emdyp* expression when compared to dormant protoscoleces, although this increase was not statistically significant. As shown in Fig. 4B, keeping metacestode vesicles in the presence of oxygen when compared to axenically cultivated vesicles, slightly increased *emdyp* expression as did *in vivo* cultivation of parasite material in the peritoneum of jirds. A highly significantly increase (more than 150-fold) of *emdyp* expression was, however, observed for metacestode tissue deriving from jird liver when compared to all other conditions. These data indicated that the parasite’s Dyp-type peroxidase encoding gene fulfils an important role within the liver during an infection of the intermediate host, probably by counteracting ROS that are produced during the host immune response.

**Fig. 4.:**
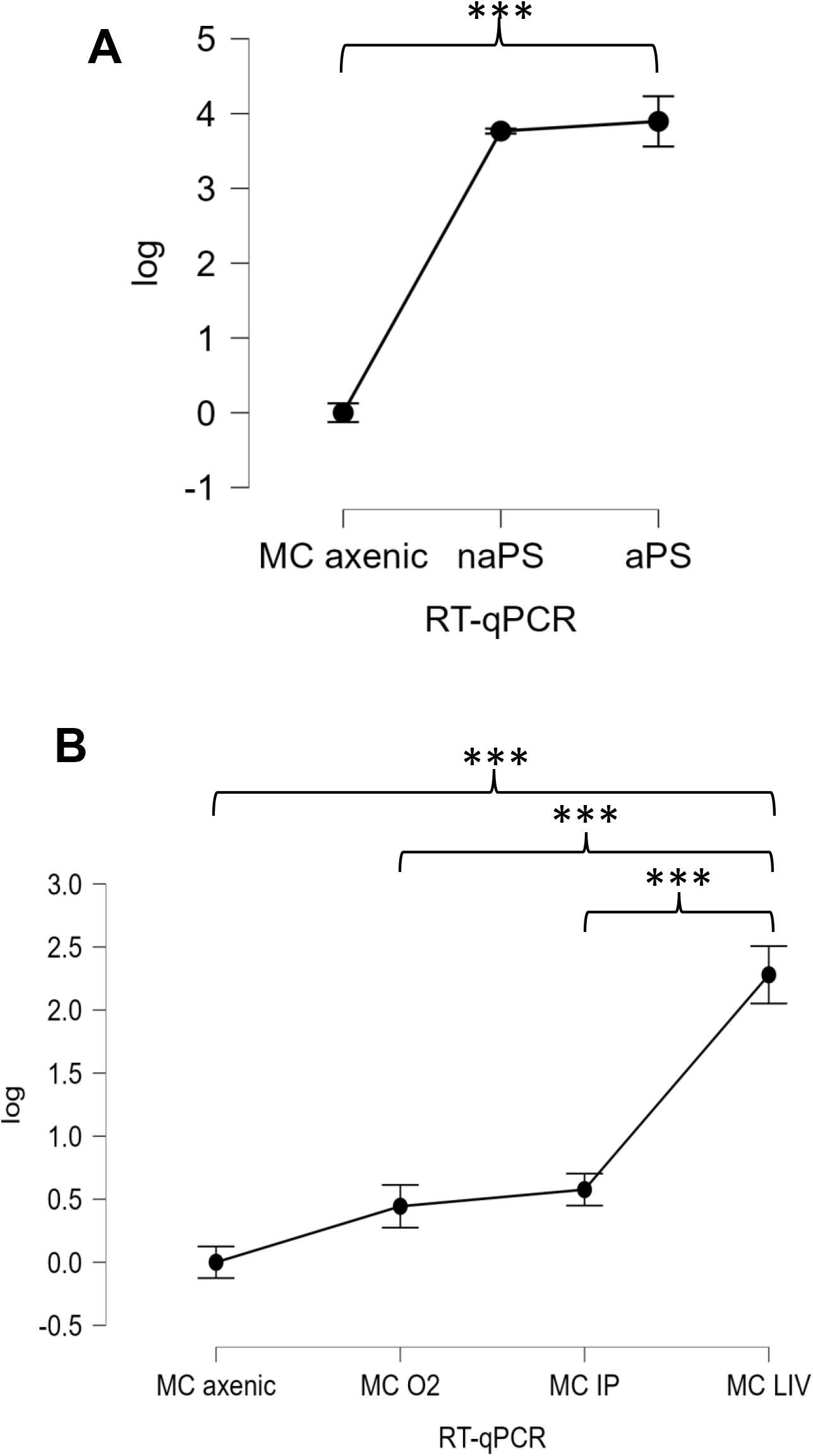
RT-qPCR on *emdyp* expression in *Echinococcus* larval stages. RT-qPCR was carried out using primers specific for *emdyp*. As a constitutively expressed control gene, *elp* (EmuJ_000485800) was used. (A) *emdyp* gene expression in non-activated (naPS) and low pH/pepsin activated protoscoleces (aPS) in comparison to axenically cultivated metacestode vesicles. (B) *emdyp* expression in metacestode tissue. Expression levels were determined for axenically cultivated vesicles (MC axenic), vesicles cultivated in the presence of feeder cells in 5% CO2 (MC O2), as well as metacestode tissue isolated from the peritoneum (MC IP) and from the liver (MC LIV) of infected jirds. Error bars represent standard deviation. Tukey’s multiple comparison test, followed by one way ANOVA was used to compare all experimental combinations. *** indicates p<0.001

## Conclusions

In the current work we cloned and characterized a Dyp-type peroxidase encoding gene of the human parasitic cestode *E. multilocularis* which, to our best knowledge, is the first gene of this type that has been functionally investigated in an animal. We demonstrate that the encoded protein, EmDyp, displays significant homologies to Dyp-type peroxidases from bacteria and fungi and contains all conserved residues of this enzyme class at the corresponding positions. Biochemical assays further demonstrated that EmDyp possesses both peroxidase and catalase activity and could, thus, play an important role in detoxifying ROS that are produced during the parasite’s aerobic metabolism as well as during the host immune response. Gene expression analyses further showed that *emdyp* is particularly highly expressed in the protoscolex, which is probably associated with its role in removing hydrogen peroxide produced during aerobic growth conditions. However, we also observed strong overe-xpression of *emdyp* within metacestode tissue isolated from infected host liver tissue. These data indicate that *emdyp*, in addition to protecting from ROS produced during aerobic growth, might also fulfil an important role in the defence against the host immune response. Since Dyp-type peroxidases are not encoded by the genomes of *Echinococcus* host species (Zamocky et al., 2015; Sugano and Yoshida, 2021), EmDyp could therefore be an attractive target for the development of small molecule compounds to treat alveolar echinococcosis. So far, Dyp-type peroxidases have exclusively been studied in the background of their industrial potential as dye decolorizing enzymes (Sugano, 2009; Colpa et al., 2013; Zamocky et al., 2015; Sugano and Yoshida, 2021) and were, thus, not subjected to screening procedures for inhibitory substances. Given that these enzymes are encoded by the genomes of a wide variety of flatworm parasites, including the genera *Echinococcus* and *Schistosoma* (Zamocky et al., 2015), we therefore propose that respective investigations should be undertaken in the future.

## Acknowledgements

This work was supported by the Wellcome Trust (https://wellcome.ac.uk/), grant 107475/Z/15/Z (FUGI, to KB), and by a grant of the Bayerische Forschungsstiftung (https://www.forschungsstiftung.de/) (AZ-1341-18) (to KB). The authors wish to thank Monika Bergmann and Dirk Radloff for excellent technical assistance. Markus Spiliotis is thanked for technical advice.

